# Evidence for Free Radical Drug Ligands in class A G-protein Coupled Receptors

**DOI:** 10.1101/2021.08.30.458164

**Authors:** Angela S Gehrckens, Andrew P Horsfield, Efthimios M C Skoulakis, Luca Turin

## Abstract

We analyse the conformation of 22 cationic ligands bound to G-protein coupled receptors of known structure and find three ligands that are present inside receptors not in their closed shell cation form, but as neutral radicals, i.e. after addition of an electron to the ligand. The implications of this finding for GPCR function are discussed and a possible experimental test is proposed.

## Introduction

Class A (rhodopsin-like) G-protein coupled receptors (GPCRs) make up more than three quarters of known mammalian GPCRs ^1^, and approximately half of Class A genes encode olfactory receptors. We have previously proposed that olfactory receptors operate by an electronic mechanism that enables them to sense odorant vibrations^2–6^. A valid objection to our proposal has been that there is so far no evidence for electron transport in GPCRs^7^. Here, using published receptor structures and density-functional energy calculations, we show that some cationic GPCR ligands in published structures are in fact present in the crystal as neutral free radicals, i.e. with an electron added to the drug molecule. We suggest this is evidence for electron movement through molecular wires within Class A GPCRs.

### Evidence for electron transfer in 5-hydroxytryptamine receptors

While examining structures of GPCR ligands, we noticed that one of them, the inverse serotonin agonist methiothepin existed in two different conformations when bound to two different 5-hydroxytryptamine receptors, 5HT-1 (pdb 5v54, 3.4Å resolution) and 5HT-2 (pdb 6wh4, 3.9 Å resolution). They are shown in figure 1. They differ in the pucker of the central thiepin and the angle between the two aromatic rings. This difference is unlikely to be the result of a poor fit to the electron density map, because the presence of two heavy sulfur atoms gives a high electron density along the bonds and constrains the conformation at the thiepin ring sulfur. The electron density score for individual atoms (EDIA) quantifies the electron density fit of each atom in a crystallographically resolved structure. EDIA scores ^8^ are shown for both structures in figure 1, showing that the position of the two benzene rings is reliably estimated from the electron density.

**Figure 1:**
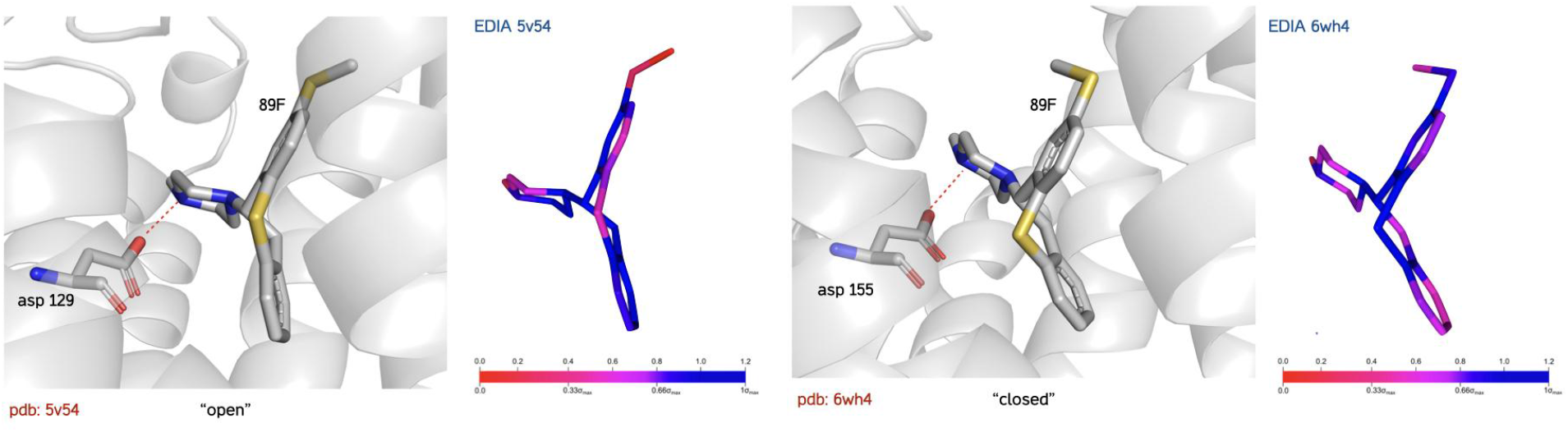
Structures of the GPCR ligand methiothepin (89F) in two different 5-HT receptors, type 1 (pdb 5v54, left) and type 2 (6wh4, right). Both ligands are protonated and form an ion pair (red dotted line) with an aspartate residue. The two methiothepin ligands are markedly different in structure, the angle between the planes of the aromatic rings is larger in the 5v54 than in the 6wh4 conformation. The left panels show the structure in the pdb file, the right panels are calculated EDIA scores for individual atoms and bonds 89F indicating the support from the electron density map for those atom positions. Blue indicates good support, red a poor one.

Methiothepin is protonated on the terminal nitrogen of the piperazine adjacent to the thiepine, and the protonated nitrogen binds to a carboxylate (asp 129 in 5v54, asp 155 in 6wh4). The conformation of methiothepin in 6wh4 is very close to the calculated closed-shell cation structure. In light of the electron transfer proposed in rhodopsin^9^, and given that the methiothepin cation contains both delocalised bonds and a positive charge, we wondered whether these two structures represented respectively the cation and the neutral free radical, i.e. whether adding an electron to the 6wh4 structure to give the neutral free radical would yield the 5v54 structure. The structure of cation and neutral free radical are shown in figure 2. The free radical is a very good match to the methiothepin structure in 5v54. The formation energy of the neutral free radical is 1.6 eV lower than that of the cation in a dielectric medium with ε=4. The energy difference between the observed structures in the absence of an added electron, is 0.25 eV, or ≈10kT which makes it unlikely but not impossible that steric constraints and binding energy alone could have deformed one into the other. Interestingly, the removal of an electron to give the dication, while energetically very unfavorable (≈6 eV), also leads to a flattened geometry.

**Figure 2.**
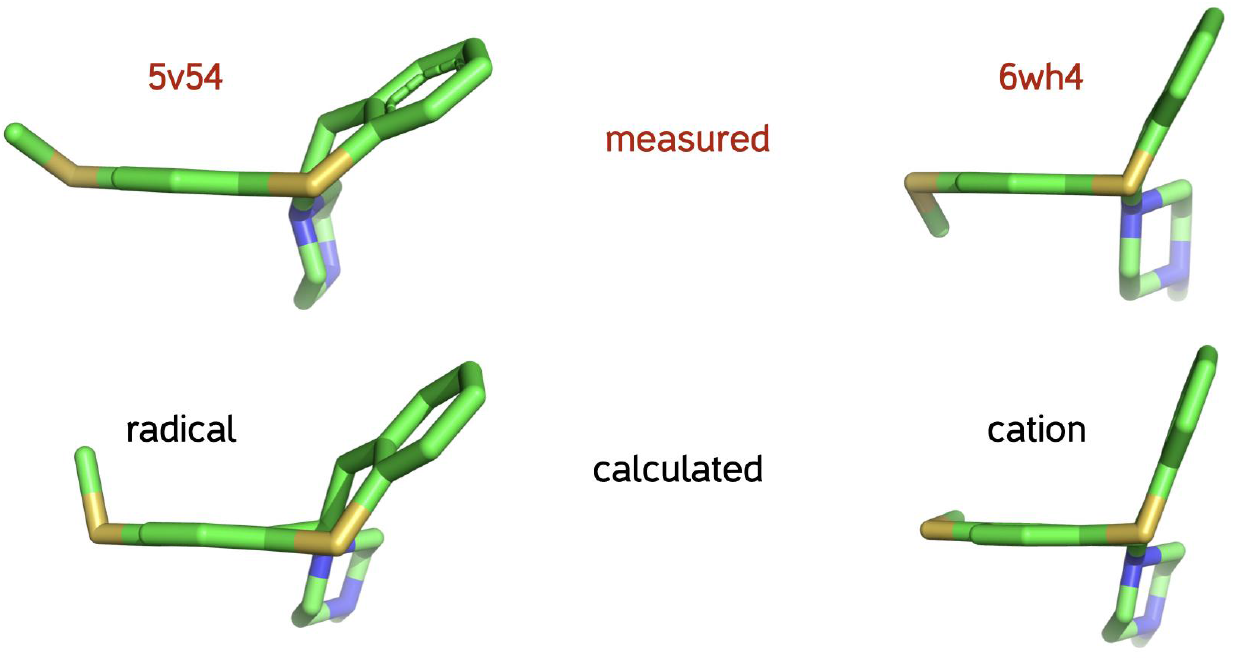
**Top**: **Left:** structure of the metitepin (pdb 89F) found in the 5HT1B receptor (pdb 5v54).**Right**: structure of the metitepin found in the 5HT1A receptor (pdb 6wh4). **Bottom: Left:** structure of the neutral free radical calculated from the native structure by protonation of the piperazine ring nitrogen adjacent to the thiepine and addition of one electron. The neutral free radical conformation closely approximates that of the pdb structure in 5v54 both in ring dihedral angle and pucker. **Right:** energy-minimised structure of the closed-shell cation, closely mimicking the structure seen in 6wh4. Structures calculated with a 6-311G* basis set with the B3LYP functional in a medium with dielectric constant ε=4 medium, using CPCM solvation.

Since the two drug conformations sit in two different receptors, it is necessary to check for the possibility that the difference in structure between the two methiothepin configurations might be due to their environment rather than electron transfer, so we calculated the conformation of bound methiothepin surrounded by its binding site amino acids. As in the previous calculations, the alpha carbons were held fixed and the structures were minimized using a dispersion-corrected hybrid functional (B3LYP-BJ) and a TZP basis set. In the absence of an added electron the charge on the methiothepin + binding site complex is zero, because the aspartate and the piperazine cancel each other. When both binding sites are calculated at zero charge, the structure in 6wh4 was essentially unaltered by minimisation, whereas the structure in 5v54 relaxed to the 6wh4 one, showing that interactions with neighboring amino acids are insufficient to maintain the structure. By contrast, if an electron is added to either methiothepin + binding site, the 5v54 conformation of methiothepin is obtained after minimisation. This agrees with the notion that the difference between the two methiothepin conformations observed in crystals is due to addition of an electron.

### Other GPCR ligands present as free radicals

It seemed interesting to ask whether other known structures of drugs bound to GPCRs show any structural evidence of electron gain to form a free radical. Structures and formation energies of closed-shell and free radical drugs can be calculated to a high accuracy using DFT, and compared to the drug structure in the crystal. There are in principle three possible cases: 1-free radical formation leads to a small structural change, in which case no determination is possible. 2-free radical formation leads to a large structural change and the drug bound to the receptor is in the closed-shell (singlet) cation conformation and 3-free radical formation gives a large structural change and the drug bound to the receptor is in the neutral free radical (doublet) conformation. It should be borne in mind that the average resolution of GPCR structures is typically of the order of 2.8Å, so that only sizable structural changes will be detected. Furthermore, in the lower-resolution structures, if the structural change is small the tendency will be to fit the electron density to the closed-shell drug conformation since crystallographers have no *a priori* reason to expect a free radical, and there may therefore be an undercount of free radical structures.

Starting with 51 ligands of Class A GPCRs reported in the RCSB database, we selected those present as cations, on the assumption that electron affinity of a neutral molecule would be insufficient to attract an electron. The presence of a protonated (cationic) nitrogen in the drug structure was ascertained by the presence in the pdb structure of a carboxylate near it. 22 such protonated structures were found (Table 1). All were calculated as both cations and neutral free radicals (B3LYP, 6-31G*) in a medium of dielectric constant ε=4. Of these, 7 displayed sizable conformational changes upon addition of an electron and were recalculated using an improved basis set (6-311G*). They were: the D2 dopamine antagonist eticlopride (chemical ID ETQ); the 5-HT agonist ergotamine (ERM); methiothepin (see above, 89F); the muscarinic antagonist quinuclidinyl benzylate (QNB); the dopamine antagonist zotepin (ZOT); and the D4 dopamine antagonist L-745,870 (L74). Of these, ETQ and QNB were unambiguously present as cations in the receptor structures. Ergotamine ERM was present as an ambiguous structure. Metitepin, Zotepin and L-745,870 were present as structures closer to the neutral free radical than the closed-shell cation. In the case of zotepin (figure 3), which has a similar structure to methiothepin and behaves similarly, the formation energy of the neutral free radical is 2.55 eV lower than that of the cation in a dielectric medium with ε=4. The difference in energy between the open and closed conformations in the absence of an added electron is 1.85 eV, which rules out both thermal motion and weak noncovalent interactions as a possible cause for its conformation *in situ*.

**Table 1:**
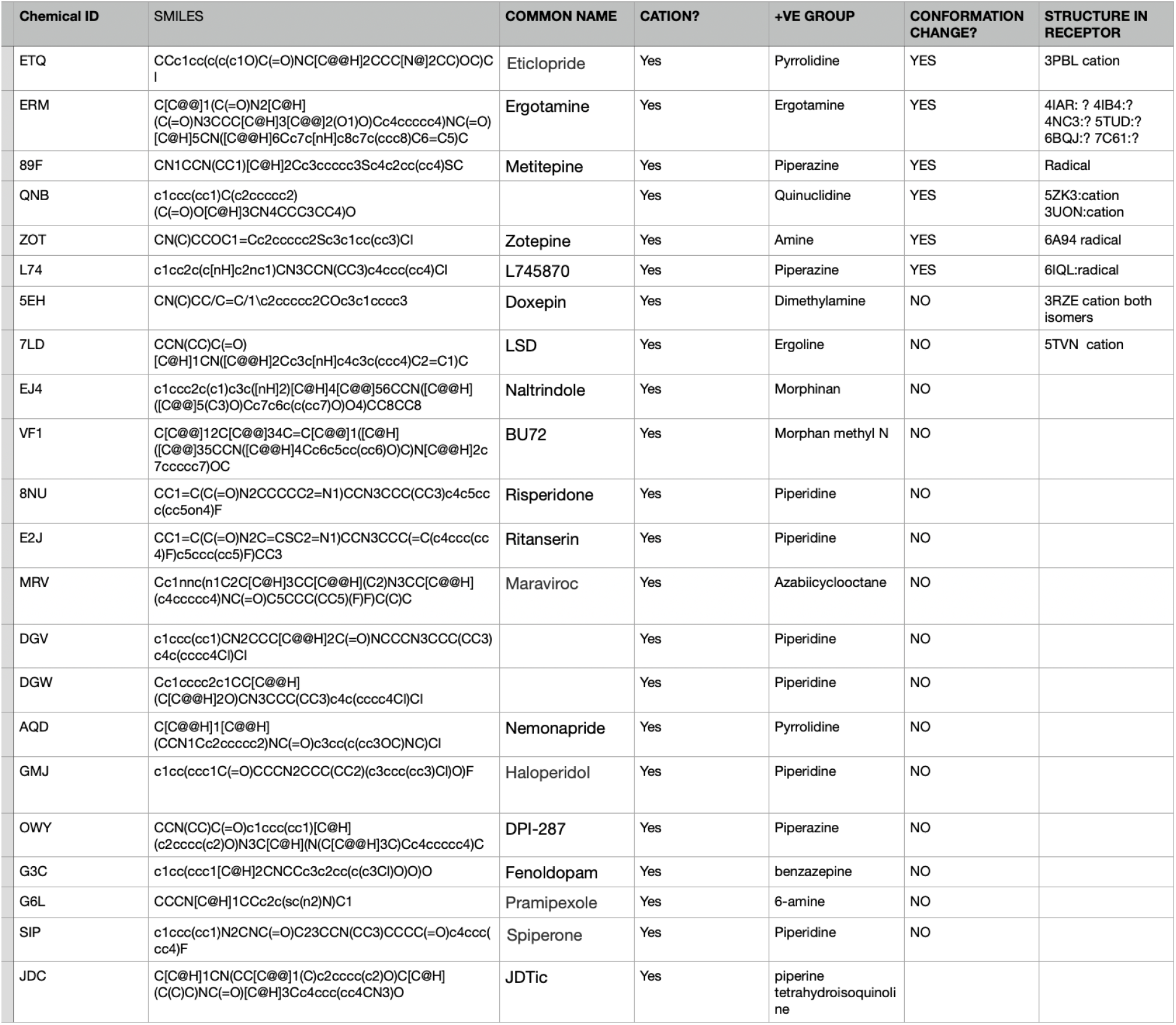
22 cationic ligands studied. Columns from left: three-letter pdb identifier, structure in Smiles notation, cationic status, common name, conformational change upon electronation and nature of the molecule in crystal.

**Figure 3.**
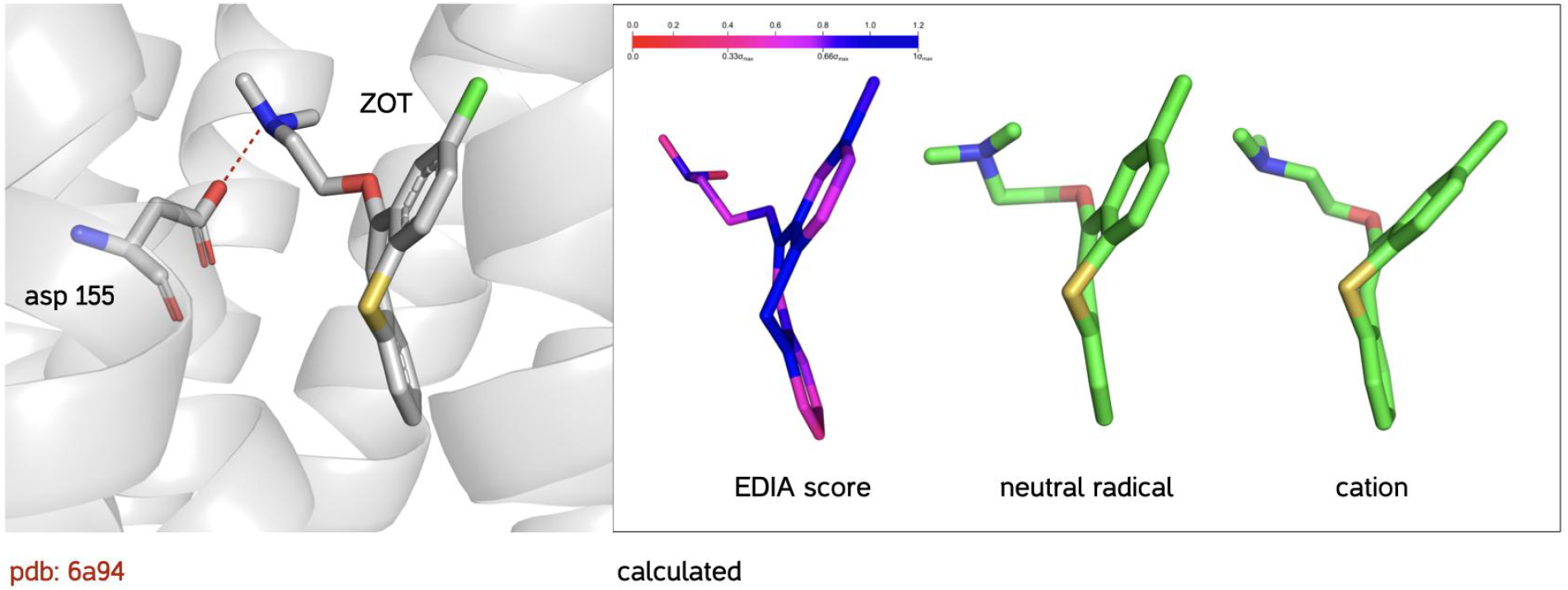
**Left:** structure of zotepin (pdb ZOT) found in the 5HT2A receptor (pdb 6a94).**Right**: EDIA score of the crystal structure; calculated structure of the neutral free radical; calculated structure of the closed-shell cation. The EDIA score shows good support for atom positions in the thiepine ring. The neutral free radical conformation closely approximates that of the pdb structure in 6a94 both in ring dihedral angle and pucker. The energy-minimised structure of the closed-shell cation resembles the structure of metitepin found in 6wh4. Structures calculated using a 6-311G* basis set with the B3LYP functional in a medium with dielectric constant ε=4, using the CPCM solvation model.

The case of L-745,870, 3-([4-(4-chlorophenyl) piperazin-1-yl]methyl) -1H-pyrrolo [2,3-b] pyridine, a Dopamine D4 subtype-selective antagonist, is less clear-cut (figure 4). The molecule has more conformational freedom than the tricyclic structures of 89F and ZOT, and the atom positions in the crystal structure have less support than for 89F and ZOT, in particular the chlorophenyl ring is relatively less well resolved. However, the main difference between the calculated free radical and cation structures is the orientation of the pyrrolopyridine (indolyl) terminal group of which three atoms have good EDIA support. The indolyl ring, on which all the spin resides, is rotated almost 180 degrees between free radical and cation, and in the absence of an added electron going from “cation” to “free radical” conformation requires 0.54 eV, or ≈20 k_B_T at 300K.

**Figure 4.**
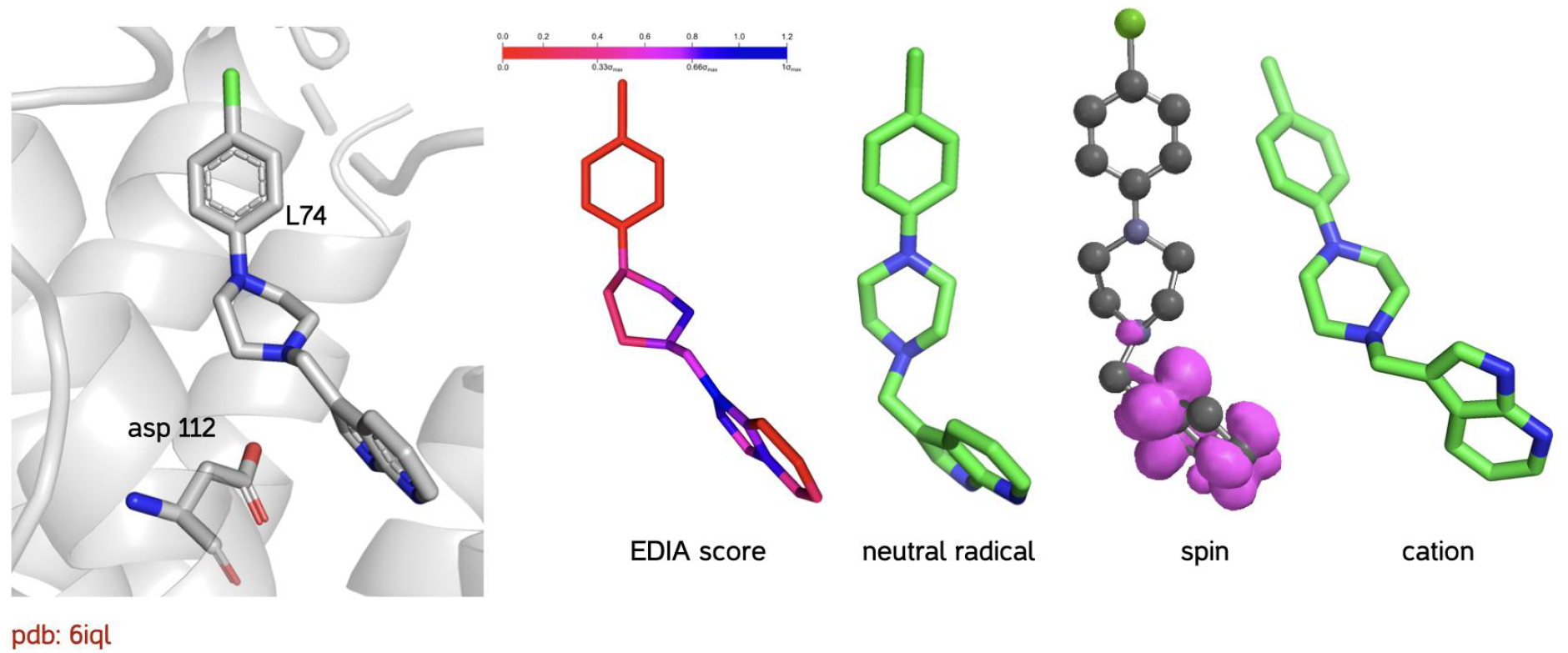
**Left:** structure of L-745,870 (pdb L74) found in the Dopamine D4 receptor (pdb 6iql).**Right**: EDIA score of the crystal structure; calculated structure of the neutral free radical; spin density surface (.002 e/au^3^) on the calculated free radical; calculated structure of the closed-shell cation. The EDIA score shows poor support for atom positions in the chlorophenyl ring. The neutral free radical conformation approximates that of the pdb structure in 6iql. The energy-minimised structure of the closed-shell cation rotates the indolyl ring by 180 degrees. Structures calculated using the 6-31G* basis set with the B3LYP functional in an implicit solvent medium with dielectric constant ε=4.

### Pathways for electron hopping within GPCRs

Considered as semiconductors, proteins have a high (≈5 eV) bandgap ^10^ and are unlikely to conduct unless doped with very strong electron-withdrawing groups ^11^. However, some amino acid side chains have favorable energetics for electron hopping. In particular, the aromatic amino acids tryptophan, phenylalanine, tyrosine and histidine have been implicated in electron and hole hopping across proteins. Hopping takes place by tunnelling, and therefore requires nanometer scale proximity between amino acids. Betts, Beratan and Onuchic^12^ have described an algorithm which enables the identification of aromatic residue wires and networks that could sustain hopping electron currents. We have used a simplified version of their algorithm called Emap https://emap.bu.edu./ ^13^, not including “through-bond” electron movement, to map potential electron wires within GPCRs. We restrict our search to the four receptors in which we have reason to suspect electron transfer is occurring: rhodopsin 1u19, the 5-HT1B receptor 5v54, 5-HT2A receptor 6wh4. Perhaps unsurprisingly given the high aromatic content of all GPCR membrane helices, all four receptors contain potential molecular electron wires spanning most or all of the membrane. They are shown in figure 5. These maps were obtained under stringent conditions of 10Å distance between aromatic amino acids. Relaxing this distance to 15 or 20 Å leads to many more paths spanning the membrane.

**Figure 5:**
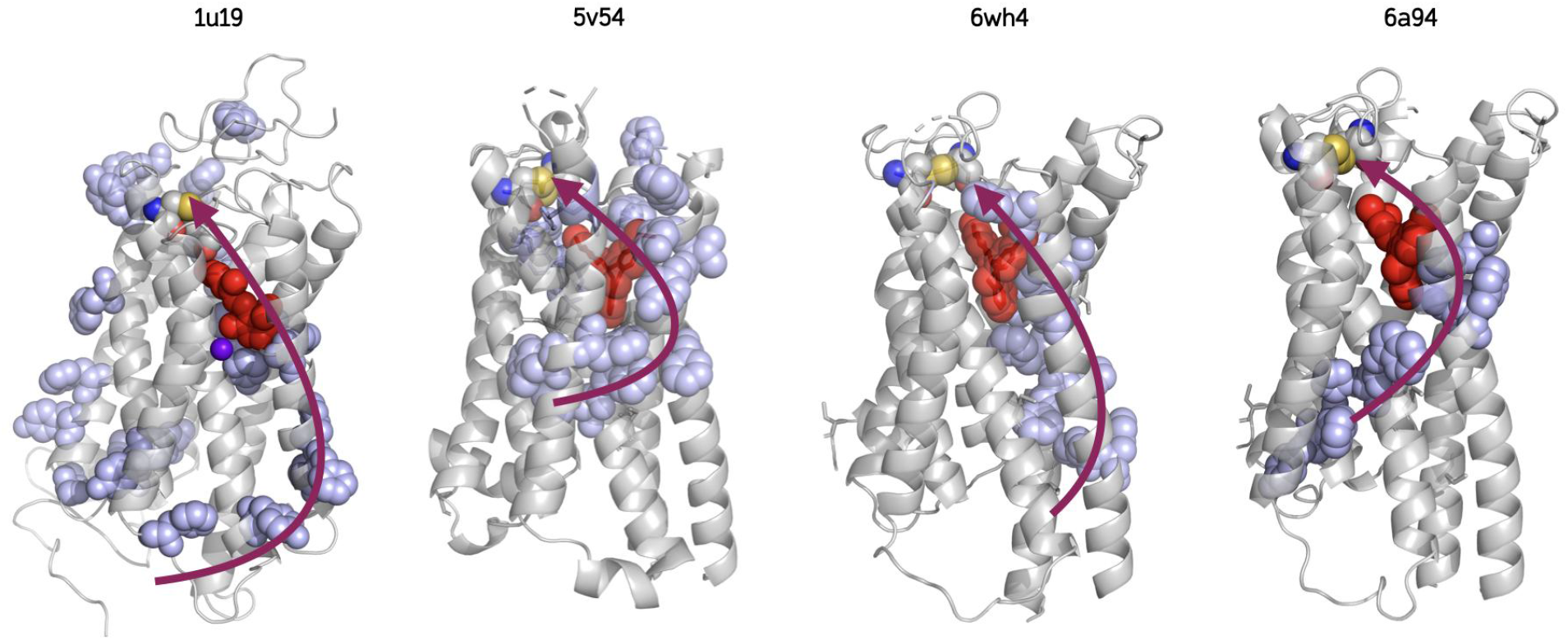
Structures of rhodopsin (1u19) and three 5-HT receptors 5v54, 6wh4 and 6a94. The side chains shown in blue are those of aromatic amino acids (trp, tyr, phe and his) situated less than 10Å from each other and therefore able to allow hopping of electrons and holes across the structure. The ligand is shown in red (retinal RET in 1u19, methiothepin (89F) in 5v54 and 6wh4, and zotepin (ZOT) in 6a94. Zinc is purple in 1u19. The sulfur atoms of the extracellular disulfide bridge present in all Class A GPCRs is shown in yellow near the ligand. Red curved arrows highlight some possible pathways for electron (or hole) transfers from the cytoplasmic face to the extracellular disulfide *via* the ligand.

**Figure 6:**
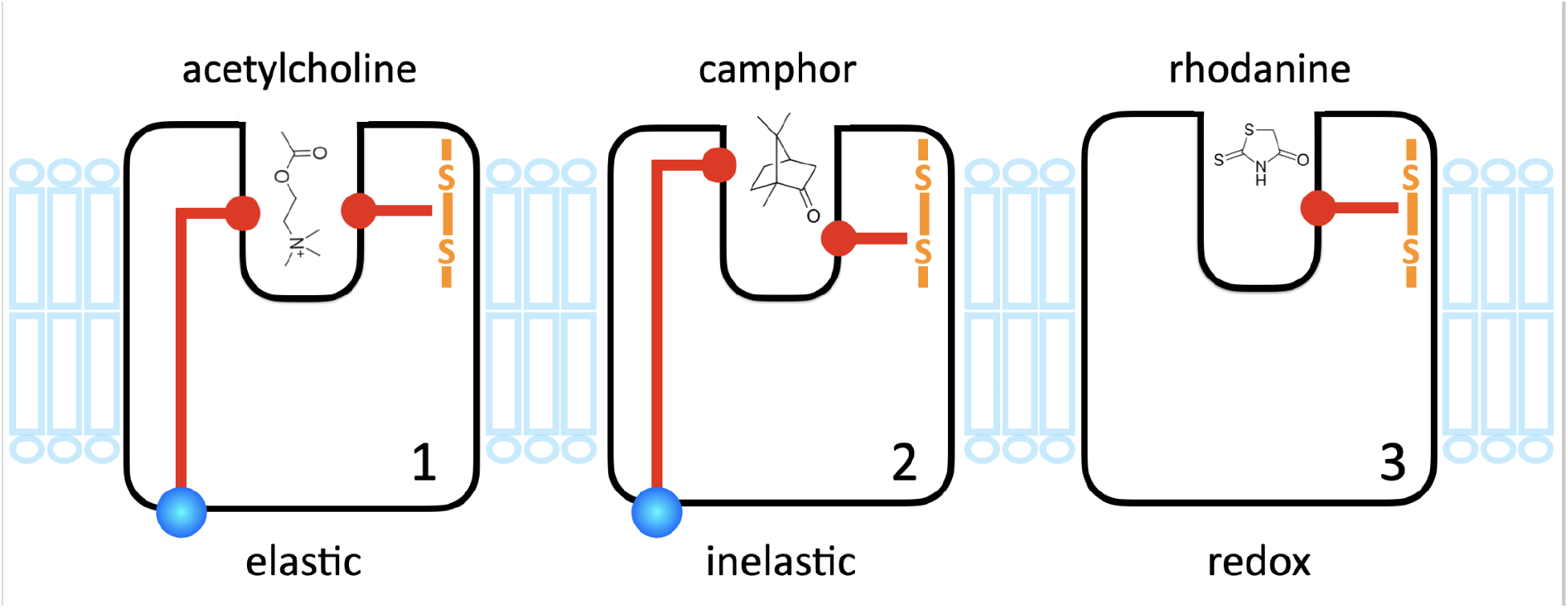
Three possible proposed schemes for an electron circuit in a GPCR. In 1, electrons flow from an intracellular binding site for soluble electron donors (blue) to an extracellular disulfide bridge, normally oxidised, via the ligand binding site, which contains a cationic drug, here acetylcholine. The positive charge facilitates elastic tunnelling across the drug to the disulfide. In 2, the donor and acceptor site on either side of the ligand lie at different energies, and inelastic tunnelling dominates, enabled by energy loss across the ligand to molecular vibrations. In 3, electron donation proceeds from the ligand itself to the disulfide bridge.

What do these structures tell us about the direction of electron flow? Again taking our reference set of GPCRs, a striking fact is that the postulated molecular wires appear able to connect the cytoplasmic face of the GPCR to the highly conserved disulfide bridge first found in rhodopsin^14–17^ close to the extracellular face *via* the bound ligand. It makes good sense for the oxidised disulfide-dithiol pair to be extracellular, since the reducing agents are inside the cell while the extracellular medium is oxidising^18,19^. The presence of an electron wire could bring electrons from cytoplasmic electron donors such as flavin nucleotides to reduce the extracellular disulfide bond, known to be essential to function^20–25^.

## Discussion

In summary, we have found structural evidence for three receptor-bound drugs being present as free radicals in two 5-HT receptors and one dopamine receptor. We have also shown that pathways for electron and hole hopping are present in the receptors of interest, potentially connecting the cytoplasmic face of the receptor to a functionally important electroactive structure, the extracellular disulfide link. The disulfide bridge which we propose as a reduction target for electron flow is highly conserved (94%) in class A GPCRs ^26^. Electron flow across plasma membrane proteins from intracellular electron donors (NADH) to the extracellular face is well documented (see ^27^ for review), though the exact mechanism remains obscure. There is also evidence of radical pair formation involving tryptophan radicals^28–30^ in the absence of light. Such “dark”radical pair formation could excite the electron which then hops from one aromatic amino acid to another.

The small number of drugs we have found that both exhibit a structural change following one-electron reduction *and* appear in the structure as radicals is not a reliable indicator of the incidence of free radical formation in Class A receptors in general. The addition of an electron to a ligand generally results in a small structural change, and therefore makes the event invisible to crystallography. However, the presence of a stable free radical in a GPCR can in principle be checked easily with an electron spin resonance spectrometer. Even basic ESR spectrometers can easily detect 10^12^ spins^31^. If each protein occupies a cube 10 nm on the side, a crystal 100µm in size should give an unequivocal signal around g=2. The crystals 5v54, 6a94 and possibly 6iql, (and their mother liquor) should be paramagnetic.

If electrons move within Class A GPCRs, what is their impact on receptor function? In rhodopsin, the presence of delocalised orbitals due to the zinc alters the light absorption spectrum and may be used in tuning receptors to different wavelengths^9^. In other GPCRs the well-documented modulation of binding and response seen with dithiol reagents^20–25^ is an indication of the functional significance of a redox pathway..

Without prejudging the ultimate effect of electron flow on receptor function, we envisage three possible mechanistic schemes. In Scheme 1 elastic tunnelling dominates. An effective way to control the rate of electron hopping is to lower the barrier to hopping using a positive charge. Charged ligands could therefore regulate electron flow. In the case of neurotransmitter receptors the ligands (e.g. catecholamines, indoleamines) are cations, as indeed are their pharmacological agonists and antagonists. The drug acts as a cationic switch for electron flow from the intracellular face (blue dot) to the extracellular disulfide bridge, normally present in the oxidised form. The term “alkaloid”, referring to natural small-molecule ligands, denotes a substance containing a basic nitrogen, typically protonated at physiological pH. Uncharged drugs are very much the exception^32^. In this view cationic drugs, once bound to their binding site by highly stereospecific interactions, may then act in the same fashion by lowering a tunneling barrier^6^.

This said, our observation that the drugs in some Class A GPCRs are present in the crystal structure as uncharged free radicals does not fit the simple scheme where a drug acts as a facilitator of electron transfer while remaining unaffected. The data is too sparse to draw any certain conclusions from this, but our findings at least suggest that electron transfer could be impeded or blocked by the presence of what is de facto a spin-trap ligand replacing the natural one. The second mechanism is a modification of the first, in which donor and acceptor levels across the ligand-binding site are shifted in energy and inelastic tunnelling dominates. This repurposes the receptor as an inelastic-tunnelling sensor which may have found a use in the detection of molecular vibrations by olfactory receptors^2–6^

Finally, many drugs are electroactive as electron donors, a property which is used in their detection in vivo^33^ and in analytical chemistry^34^. There may be situations where the drug itself is a strong enough electron donor to reduce the disulfide bridge without external electron input. This is depicted in scheme 3 (redox). This mechanism may also explain some of the known pan-assay interference compounds (PAINS). ^35–37^. For example the rhodanine moiety, a well-known PAIN,^36^ is a redox-active agent^38^.

## Methods

DFT calculations were done using the Amsterdam Density Functional code^39^ (ADF, www.scm.com) version 2019.102 and later, running either on a 6-core MacBook Pro, on 50-120 cores at www.crunchyard.com or on a 64-core Lenovo P620. Ligand structures were computed using Spartan 18 (www.wavefun.com) using its molecular alignment tool to compare structures. Parameters are indicated where appropriate. Protein model alignments were made using Yasara^40^ www.yasara.com. Structures were visualised with PyMol ^41^ and Chimera^42^. Data were plotted using IgorPro (www.wavemetrics.com). EDIA scores were computed with https://tinyurl.com/49tfj99a, and electron pathways with https://emap.bu.edu/ using all-aromatics, closest atom, 10Å distance settings.

## Acknowledgments

LT wishes to thank David Briggs for advice on the interpretation of structural data. This work was supported by the Stavros Niarchos Foundation and Ionis Pharma (LT), and an Erasmus+ scholarship to AG. Data described in the present article will be supplied on reasonable request.

